# *Saccharomyces cerevisiae* Ecm2 Modulates the Catalytic Steps of pre-mRNA Splicing

**DOI:** 10.1101/2020.09.05.284281

**Authors:** Clarisse van der Feltz, Brandon Nikolai, Charles Schneider, Joshua C. Paulson, Xingyang Fu, Aaron A. Hoskins

## Abstract

Genetic, biochemical, and structural studies have elucidated the molecular basis for spliceosome catalysis. Splicing is RNA catalyzed and the essential snRNA and protein factors are well-conserved. However, little is known about how non-essential components of the spliceosome contribute to the reaction and modulate the activities of the fundamental core machinery. Ecm2 is a non-essential yeast splicing factor that is a member of the Prp19-related complex of proteins. Cryo-electron microscopy (cryo-EM) structures have revealed that Ecm2 binds the U6 snRNA and is entangled with Cwc2, another non-essential factor that promotes a catalytically active conformation of the spliceosome. These structures also indicate that Ecm2 and the U2 snRNA likely form a transient interaction during 5’ splice site (SS) cleavage. We have characterized genetic interactions between ECM2 and alleles of splicing factors that alter the catalytic steps in splicing. In addition, we have studied how loss of ECM2 impacts splicing of pre-mRNAs containing non-consensus or competing SS. Our results show that ECM2 functions during the catalytic stages of splicing. It facilitates the formation and stabilization of the 1^st^-step catalytic site, promotes 2^nd^-step catalysis, and permits alternate 5’ SS usage. We propose that Cwc2 and Ecm2 can each fine-tune the spliceosome active site in unique ways. Their interaction network may act as a conduit through which splicing of certain pre-mRNAs, such as those containing weak or alternate splice sites, can be regulated.

## INTRODUCTION

Spliceosomes are composed of small ribonucleoprotein particles (snRNPs), each containing proteins and a small nuclear RNA (U1, U2, U4, U5, or U6 snRNA), and dozens of additional protein splicing factors. Spliceosomes assemble from these factors before undergoing a number of conformational changes to form a catalytic center (activation) capable of carrying out the chemical steps of splicing (**Fig. 1A**): 5′ splice site (SS) cleavage (1^st^ step) and exon ligation (2^nd^ step). Significant genetic, biochemical, and structural work over the past few decades has provided a wealth of information into how essential components of the splicing reaction such as the snRNAs, Prp8 protein, and DExD/H-box ATPases promote splicing (Wahl et al. 2009; Yan et al. 2019; Plaschka et al. 2019; Kastner et al. 2019; Mayerle and Guthrie 2017). In comparison, much less is known about how non-essential factors modulate the splicing reaction and interact with the core machinery.

**Figure 1.**
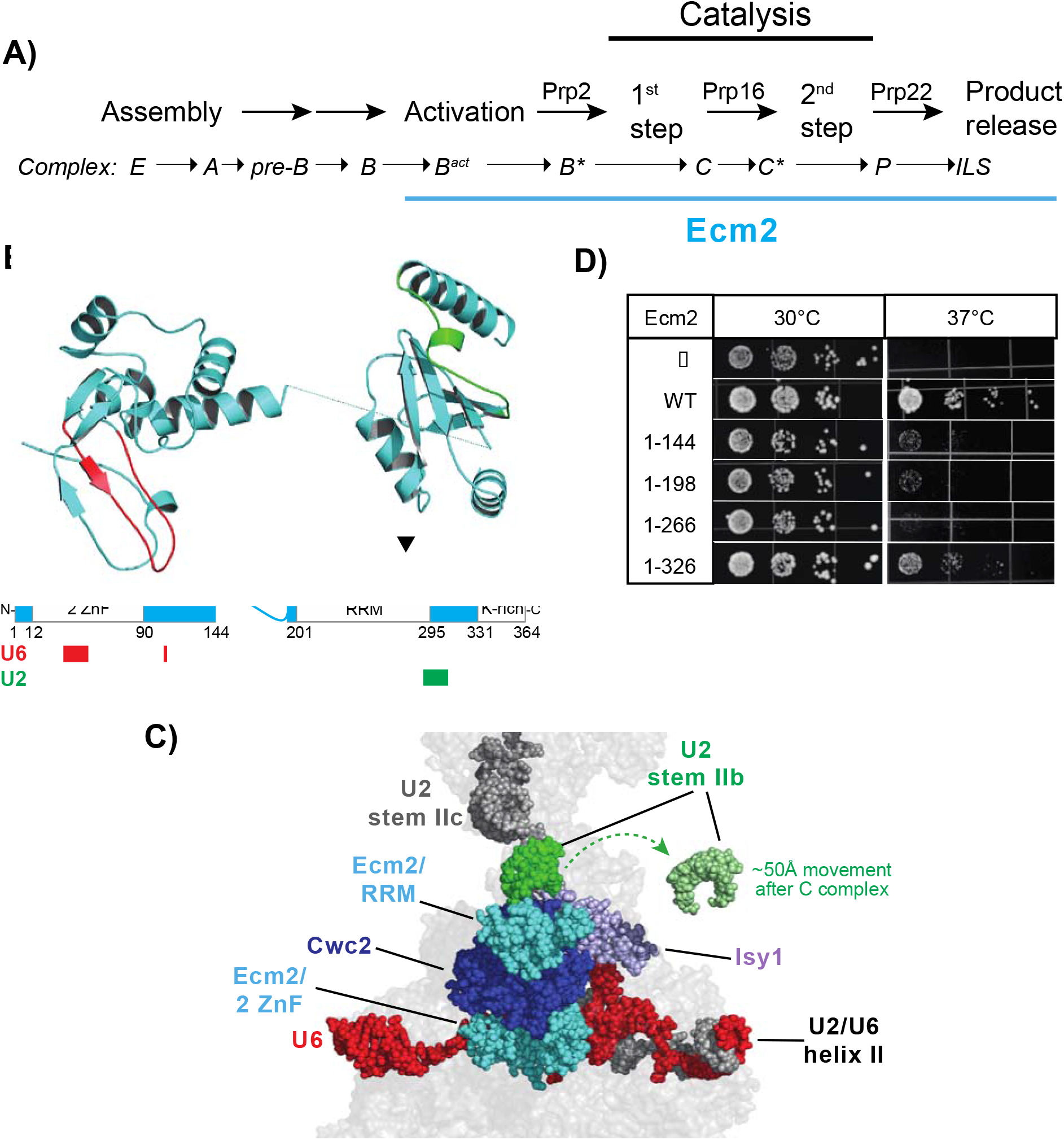
Structural Analysis of Ecm2 during Splicing. (**A**) Schematic of the pre-mRNA splicing pathway. ATPases tested for genetic interactions with Ecm2 are shown above the arrows of the respective steps that they promote. Spliceosome complexes containing Ecm2 are underlined in blue. (**B**) Cryo-EM structure and domain organization of Ecm2. U6 and U2 snRNA interacting regions are colored in red and green, respectively. Locations of truncation mutants studied in panel (C) are indicated. Structure from 6EXN.pdb. (**C**) Structure of the Cwc2/Ecm2/Isy1 hub and Ecm2-RRM/U2 stem II interaction in C complex. The position of stem IIb after remodeling in C* complex has been superimposed on this structure. The U6 snRNA, Cwc2 and Ecm2 do not significantly change positions in C* complex. This figure was created using 5LJ5.pdb and 5MQ0.pdb. (**D**) Temperature sensitivity of *ecm2Δ* and truncation mutants on-ura dropout media after 3 days of growth.

Several of the non-essential splicing factors in yeast are associated with the Prp19-containing complex (NTC). Indeed, of 26 yeast proteins categorized as core-NTC components or NTC-associated, 12 are non-essential (Hogg et al. 2010, 2014). Despite not being critical for growth, many of these proteins are well-conserved and have human splicing factor homologs. In general, the NTC is thought to stabilize catalytic conformations of the U6 snRNA and contribute to splicing fidelity (Hogg et al. 2010). Consistent with this model, cryo-EM structures and biochemical assays have shown that several non-essential NTC proteins directly interact with U6 including Cwc2, Ecm2, and Isy1 (Plaschka et al. 2019; Hogg et al. 2010; McGrail et al. 2009; Villa and Guthrie 2005; Rasche et al. 2012). Exactly how the NTC modulates RNA interactions within the spliceosome is not yet clear (Hogg et al. 2010), and it is difficult to infer potential mechanisms and the impact of mutations from cryo-EM structures alone (Mayerle and Guthrie 2017).

The NTC-associated protein Ecm2 was first isolated in a synthetic lethality screen for genetic interactors with U2 snRNA mutations (synthetic lethality with U2/*slt* screen) (Xu et al. 1998). This screen identified *slt11*/*ecm2* as well as other splicing factors including Prp8 and Brr2. Additional work demonstrated genetic interactions between *ecm2* and multiple components of the spliceosome but especially U2/U6 helix I and helix II mutants (Xu and Friesen 2001; Xu et al. 1998). Biochemical assays of splicing and spliceosome assembly showed that absence of Ecm2 results in loss of splicing activity at high temperatures and a block in spliceosome activation (Xu and Friesen 2001). Since it was known that U2/U6 form an intermolecular duplex during the early stages of activation (helix II) (Wassarman and Steitz 1992), it was proposed that Ecm2 functions during spliceosome activation to facilitate formation this duplex (Xu and Friesen 2001).

Structures of Ecm2 integrated into a number of spliceosome complexes have been determined by cryo-EM (Plaschka et al. 2019). Ecm2 contains two RNA binding domains separated by a linker: an N-terminal zinc finger motifs (ZNF) domain and a C-terminal RNA recognition motif (RRM) (**Fig. 1B**). Unexpectedly, Ecm2 does not directly bind U2/U6 helix II. Rather the ZNF domain interacts with the U6 nucleotides (nt) 29-32, which are located between the U6 5′ stem loop and the ACAGAGA-box/5′ SS pairing region (**Fig. 1C**). Ecm2 is also intertwined with another NTC-associated protein, Cwc2. Cwc2 contacts both the intronic RNA downstream of the 5′ SS and the U6 snRNA at multiple locations.

Interestingly, the C-terminal RRM of Ecm2 closely approaches U2 snRNA stem IIb in the C complex spliceosome (**Fig. 1C**). Due to low resolution within this region, the exact molecular contacts between the RRM, U2 stem IIb, and nearby regions of Cwc2 are unclear. This interaction is likely transient: the Ecm2/U2 interaction has only been observed in structures captured just before (B* complex) and after (C complex) the 1^st^ step of splicing (Galej et al. 2016; Wan et al. 2019). Large conformational changes place U2 stem IIb far away from any possible Ecm2 interaction in other structures (**Fig. 1A, C**) (Rauhut et al. 2016; Yan et al. 2016a, 2016b; Fica et al. 2017). Toggling of Ecm2/U2 stem IIb contacts on-or-off in different complexes resembles structural toggle switches reported for other splicing factors. These include the RNaseH domain of Prp8, the U4/U6 di-snRNA, as well as interconversion of U2 stem II itself between two mutually exclusive structures: stem IIa/b and stem IIb/c (Perriman and Ares 2010, 2007; Hilliker et al. 2007; Mayerle et al. 2017; Abelson 2017; Rodgers et al. 2015, 2016). The significance of the Ecm2/stem IIb interaction has not been studied.

Human spliceosomes do not contain direct homologs of Cwc2 and Ecm2. Instead, a single protein, RBM22, binds the U6 snRNA at the corresponding positions. Based on limited sequence homology, it has previously been proposed that RBM22 represents a fusion of Cwc2 and Ecm2 (Rasche et al. 2012). This is supported by biochemical studies that show a similar function for Cwc2 and RBM22 in stabilizing the spliceosome active site (Hogg et al. 2014; McGrail et al. 2009; Rasche et al. 2012). In addition, both Cwc2 and RBM22 interact with intronic RNA at a location downstream of the 5′SS (Kastner et al. 2019; Rasche et al. 2012). Whether or not Ecm2 and RBM22 also share any conserved functions is unknown.

We have studied the genetic interactions between Ecm2 and splicing factors capable of modulating the 1^st^ and 2^nd^ steps of splicing including the Prp2 and Prp16 ATPases, Prp8, and U6 snRNA. Ecm2 exhibits genetic interactions with mutations that disrupt U2 stem II toggling, consistent with a functional interaction between the protein and U2. Genetic deletion of ECM2 changes how pre-mRNAs containing non-consensus splice sites are processed, implicating Ecm2 in the catalytic steps of splicing in addition to its role in activation. Our results support a model in which Ecm2 has distinct functions for each catalytic step and are consistent with a proposal that several non-essential splicing factors (Ecm2, Cwc2, Isy1) function as a hub for regulating spliceosome catalysis (Hogg et al. 2010). These results have implications for the function of RBM22 in human spliceosomes as well as for how RBM22/intronic RNA interactions are formed.

## RESULTS

### The Ecm2 U6-Binding Domain is Insufficient to Rescue Yeast Growth at 37°C

Previous studies of Ecm2 reported that *ecm2Δ* yeast exhibited a strong temperature-sensitive (*ts*) phenotype with significantly reduced or no growth at temperatures above 33°C (Xu and Friesen 2001). We replicated this result by deleting *ECM2* from a haploid strain of yeast and introducing plasmids containing *ecm2* variants under control of their endogenous promoters. As expected, *ecm2Δ* yeast containing an empty plasmid grew well at permissive temperatures (16-30°C) but possessed a severe *ts* phenotype at 37°C (**Fig. 1D**). When we included a plasmid containing the wild type (WT) *ECM2* gene, the *ts* phenotype was corrected, and growth was restored at 37°C.

To test if the N-terminal, U6-binding binding domain of Ecm2 alone was capable of rescuing the *ts* phenotype, we used recent cryo-EM structures of yeast spliceosomes to design truncation mutants of Ecm2. Nonsense mutations were incorporated at amino acids 144, 198, 266, and 326 to allow for expression of variants containing only the U6-binding ZNF domain (Ecm2^1-143^), the ZNF domain plus the inter-domain linker (Ecm2^1-197^), the ZNF and a partial U2-binding, RRM domain (lacking amino acids that come nearest to U2, Ecm2^1-265^), or the complete ZNF and RRM domains truncated at the last amino acid modeled into cryo-EM density but missing the C-terminal lysine-rich region (Ecm2^1-325^, **Fig. 1D**). Variants containing the U6-binding, ZNF domain but not the RRM were able to partially rescue the *ts* phenotype but still grew poorly at 37°C. Inclusion of the entire U2-binding, RRM (Ecm2^1-325^) resulted in more significant suppression of the *ts* phenotype; although, cells still grew more slowly than those containing Ecm2^WT^. These data are consistent with the U6-binding, ZNF domain alone being unable to completely restore Ecm2 function and the U2-binding, RRM domain contributing to this function.

### Genetic Interactions between Ecm2 and the Prp2 and Prp16 ATPases

Xu and Friesen provided ample evidence that Ecm2 plays a role in spliceosome activation (Xu and Friesen 2001). We and others have previously noted that key players in the activation process such as the U2 snRNP protein Hsh155/SF3B1 and U2/U6 helix I exhibit genetic interactions with a cold-sensitive (*cs*) mutant of the DEAH-box ATPase Prp2 (Prp2^Q548N^) (Kaur et al. 2020; Carrocci et al. 2017; Wlodaver and Staley 2014). Prp2 binds the intronic RNA downstream of the branch site and uses ATP hydrolysis to trigger release of Hsh155/SF3B1 and other U2 snRNP proteins during activation (Lardelli et al. 2010; van der Feltz and Hoskins 2019). We tested if *ecm2Δ* would also show a genetic interaction with Prp2^Q548N^. When we combined Prp2^Q548N^ with *ecm2Δ*, we observed no growth at low or high temperatures (16, 23, or 37°C) and reduced growth at 30°C (**Fig. 2A**). Prp2^Q548N^ is synthetic lethal with *ecm2Δ* at low temperatures and Prp2^Q548N^ does not rescue the *ts* phenotype of *ecm2Δ*. This genetic interaction is consistent with Ecm2’s function in promoting spliceosome activation.

**Figure 2.**
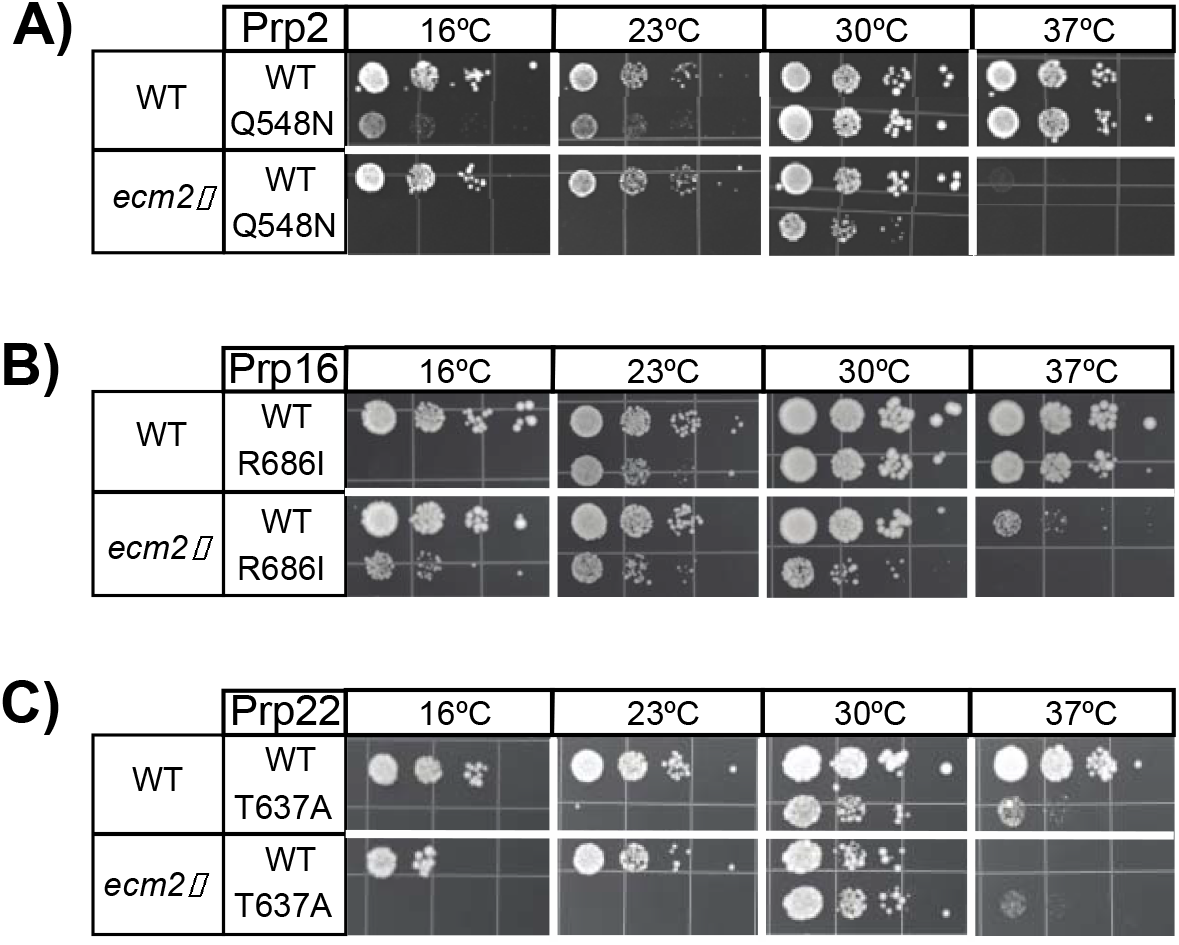
Genetic Interactions between Ecm2 and Spliceosomal ATPases. (**A-C**) Mutations in Prp2 (A, *cs*), Prp16 (B, *cs*), and Prp22 (C, *cs* and *ts*) were combined with *ecm2Δ* and tested for suppression or exacerbation of temperature-dependent growth phenotypes. Yeast were plated on YPD media and imaged after 3 (23, 30, or 37°C) or 10 (16°C) days of growth.

We next tested if other spliceosome DEAH-box ATPases would also show genetic interactions with *ecm2Δ* or if these results were specific to Prp2^Q548N^. We combined *ecm2Δ* with a *cs* mutation of the ATPase Prp16 (Prp16^R686I^) or a *cs* and *ts* mutation of the ATPase Prp22 (Prp22^T637A^). Prp16 uses ATP hydrolysis to promote conformational changes of the spliceosome and splicing factor release during remodeling of the active site from the 1^st^ to 2^nd^ catalytic step (**Fig. 1A**) (Semlow et al. 2016; Plaschka et al. 2019; Schwer and Guthrie 1992). Prp16^R686I^ likely impedes this transition since this mutation is rescued by alleles of Prp8 that promote exon ligation (2^nd^-step alleles, discussed below) (Query and Konarska 2004). Prp22 also uses ATP hydrolysis to promote conformational change that enables release of the spliced mRNA product from the active site (**Fig. 1A**) (Semlow et al. 2016; Plaschka et al. 2019; Schwer 2008; Wagner et al. 1998). In this case, Prp22^T637A^ likely impedes mRNA release and transition of the active site out of the exon ligation conformation since this mutation is exacerbated by 2^nd^-step alleles of Prp8 (Query and Konarska 2012).

When Prp16^R686I^ was combined with *ecm2Δ*, the *cs* phenotype of Prp16^R686I^ was suppressed and growth was restored at 16°C (**Fig. 2B**). Yeast containing both Prp16^R686I^ and *ecm2Δ* also grew at 23 and 30°C, albeit less well than when WT alleles were present. In addition, Prp16^R686I^ exacerbated the *ts* phenotype at 37°C of *ecm2Δ* yeast. This indicates some degree of synthetic lethality between *ECM2* and *PRP16* at high temperatures and is consistent a previous report of synthetic lethality between the *slt11-1* and *prp16-1* alleles (Xu et al. 1998). On the other hand, the *cs* phenotype of Prp22^T637A^ was not suppressed by deletion of *ecm2* (**Fig. 2C**). Yeast containing both Prp22^T637A^ and *ecm2Δ* grew very poorly at 37°C, and it was difficult to determine if Prp22^T637A^ was a weak suppressor of the *ts* phenotype of *ecm2Δ* yeast. In sum, these data strongly support genetic interactions between *ecm2Δ* and the Prp2 and Prp16 ATPases. Loss of Ecm2 exacerbates a *cs* defect in spliceosome activation caused by Prp2^Q548N^ and suppresses a *cs* defect in the 1^st^-to-2^nd^ step conformational change caused by Prp16^R686I^.

### Genetic Interactions between Ecm2 and Mutations in U2 snRNA Stem II

The above results are consistent with a model in which Ecm2 stabilizes the 1^st^-step conformation of the spliceosome: aiding its formation during Prp2-initiated activation and inhibiting its remodeling by Prp16. To gain further insight into Ecm2’s role during these steps, we combined *ecm2Δ* with mutations in the stem II region of the U2 snRNA which undergo a conformational change during activation. This region of U2 includes stem IIa/c as well as stem IIb—the RNA contacted by the C-terminal RRM of Ecm2 in cryo-EM structures of B* and C complex spliceosomes (**Fig. 1C**).

During activation, stem II undergoes a reversible conformational change from the stem IIa to the stem IIc structure, while stem IIb remains intact (**Fig. 3A**) (van der Feltz and Hoskins 2019). 5′ SS cleavage is inhibited when formation of stem IIc is blocked by deletion of the 3′ stem (ΔCC) or destabilized by mutation (Hilliker et al. 2007; Perriman and Ares 2007). In contrast, stabilization of stem IIc with additional base pairs (IIc+) promotes the 1^st^ step of splicing (Perriman and Ares 2007). Like *ecm2Δ*, mutations that destabilize stem IIc or disrupt an interaction that is physically mutually exclusive with stem IIa also suppress Prp16 mutants defective in remodeling the 1^st^-step spliceosome active site (Hilliker et al. 2007; Perriman and Ares 2007). We predicted that if Ecm2 is facilitating activation by assisting stem IIc formation, then deletion of *ECM2* should exacerbate the phenotypes of mutants that antagonize stem IIc.

**Figure 3.**
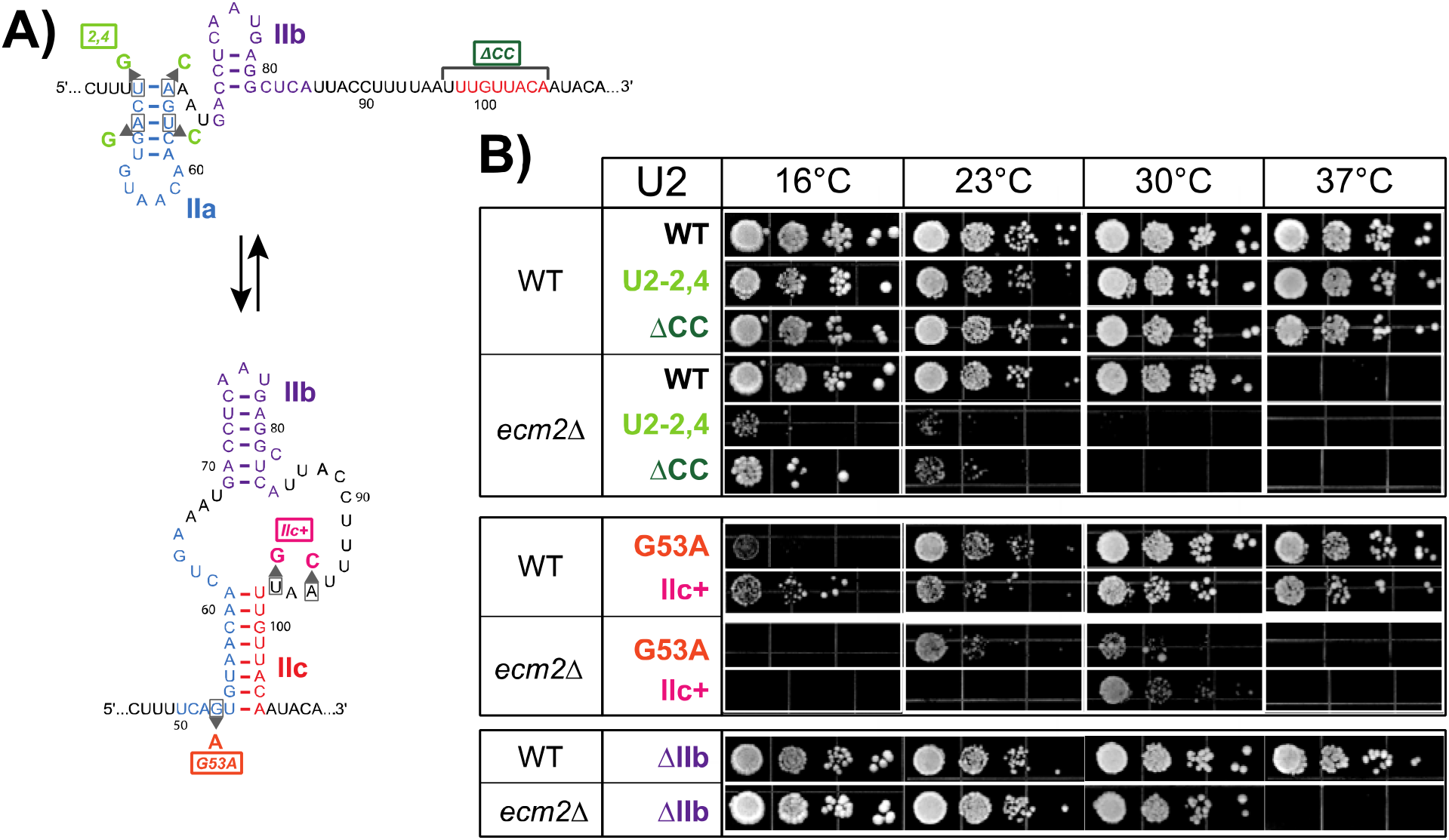
Genetic Interactions between Ecm2 and U2 stem II Mutations. (**A**) Schematic of stem IIa/IIc toggling. Mutations which disfavor stem IIc (U2-2,4 and ΔCC’. green-colored labels) are shown in the stem IIa structure. Mutations which disfavor stem IIa (G53A, IIc+; red-colored labels) are shown in the stem IIc structure. Nucleotides that are deleted in the ΔIIb mutant are colored in purple. (**B**) Mutations in stem II were combined with *ecm2Δ* and tested for suppression or exacerbation of temperature-dependent growth phenotypes. Yeast were plated on YPD media and imaged after 3 (23, 30, or 37°C) or 10 (16°C) days of growth.

The U2-2,4 and ΔCC mutations both disrupt stem IIc formation: U2-2,4 stabilizes the competing stem IIa structure while ΔCC prevents stem IIc formation entirely by deletion of the nucleotides that comprise the 3′ half of stem IIc (**Fig. 3A**) (Perriman and Ares 2007). These mutations have little phenotypic effect by themselves in our assay. However, when combined with *ecm2Δ* these mutations caused synthetic lethality at 30°C and *cs* phenotypes at 16 and 23°C (**Fig. 3B**). These results agree with our prediction that Ecm2 facilitates stem IIc formation.

This model also predicts that mutations in stem II that promote stem IIc formation may be able to suppress the *ts* phenotype of *ecm2Δ* yeast. The G53A and IIc+ mutants both favor stem IIc: G53A destabilizes the competing stem IIa structure while IIc+ extends base pairing of IIc (**Fig. 3A**) (Perriman and Ares 2007). These U2 mutants exhibit phenotypes even in the presence of Ecm2: both are *cs* while IIc+ also exhibits a modest growth defect at 30 and 37°C. Neither mutation suppressed the *ts* phenotype of *ecm2Δ* yeast, and *ecm2Δ* exacerbated the *cs* phenotypes of both mutations. These latter results could mean that Ecm2 has additional functions in the spliceosome while stem IIa is present or that snRNA structures containing these mutations are also disruptive for growth at lower temperatures in the absence of Ecm2.

If Ecm2 functions to assist stem IIc formation during activation, it is possible that this occurs through capture of stem IIb by the C-terminal RRM of Ecm2 during the B^act^ to B* complex transition. Stem IIb is non-essential in yeast (Ares and Igel 1990), and we tested if deletion of stem IIb (ΔIIb) resulted in a similar *ts* phenotype as *ecm2Δ*. The ΔIIb mutant yeast were not *ts* and exhibited minimal or no temperature-dependent phenotypes (**Fig. 3B**). The *ts* phenotype at 37°C of *ecm2Δ* was still observed when combined with the U2 ΔIIb mutation, and yeast containing both *ecm2Δ* and U2 ΔIIb grew similarly at other temperatures. This indicates that disruption of the Ecm2-RRM/stem IIb interaction is not solely responsible for the *ts* phenotype in *ecm2Δ* yeast.

### Ecm2 Impacts Splicing of Reporter pre-mRNAs Containing Non-consensus SS

We next studied how Ecm2 influences splicing catalysis *in vivo* with the ACT1-CUP1 splicing reporter (**Fig. 4A**). In this assay, splicing of the ACT1-CUP1 pre-mRNA is necessary for growth of a Cu^2+^ sensitive yeast strain on Cu^2+^-containing media. The highest [Cu^2+^] at which growth is observed is proportional to the amount of spliced mRNA in the cell (Lesser and Guthrie 1993). In the presence of an ACT1-CUP1 reporter containing consensus SS, we observed no difference in Cu^2+^ tolerance between strains with or without *ECM2*. We used a primer extension assay to confirm that the similar Cu^2+^ tolerance results were correlated with high splicing efficiencies for both catalytic steps in the presence or absence of Ecm2 (**Supplemental Fig. S1**). This indicates that splicing can still occur efficiently in the absence of Ecm2 and is consistent with lack of a significant growth phenotype in *ecm2Δ* strains at 30°C in our assays.

**Figure 4.**
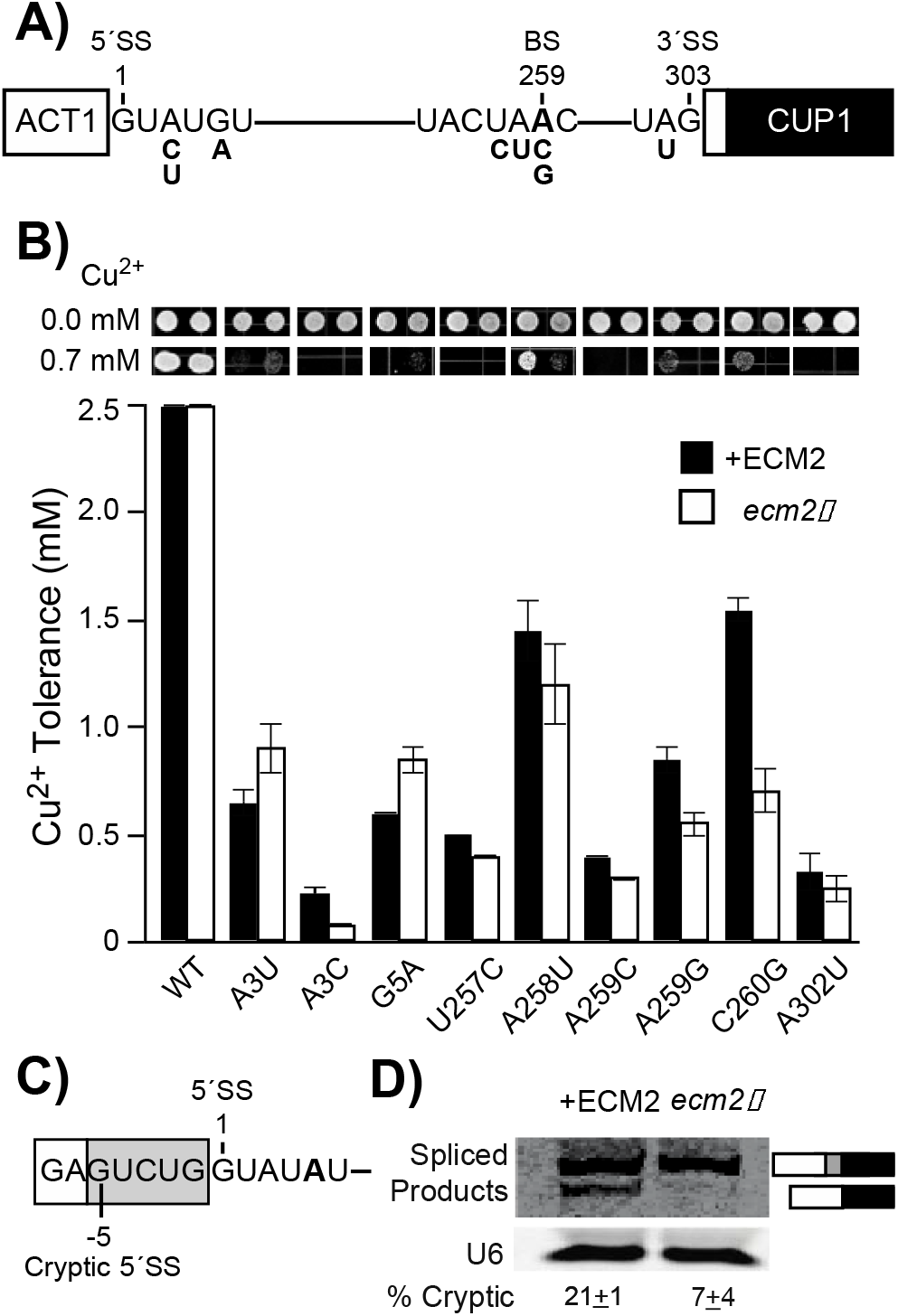
Impact of Ecm2 on Splicing of ACT1-CUP1 Reporter pre-mRNAs. (**A**) Schematic of the ACT1-CUP1 reporter pre-mRNA with non-consensus substitutions noted. (**B**) ACT1-CUP1 assay results. Representative images of yeast growth after 3 days at 30°C on agar plates made with-leu-trp dropout media containing 0 or 0.7 mM Cu^2+^ are shown above the bar graph. Each value in the graph represents the average of the highest concentration of Cu^2+^ at which growth was observed in at least three replicate assays. Error bars represent the standard deviation. (**C**) Schematic of the modified ACT1-CUP1 reporter containing a competing, cryptic 5′ SS. (**D**) Primer extension assay of cryptic 5′ SS usage using the reporter shown in panel (C). Primer extension of the U6 snRNA was included as a control. The percentages of cryptic products (ratios of cryptic products/total products) are shown below the gel and are the averages of three replicate experiments ± the standard deviation.

We then assayed splicing in *ecm2Δ* yeast using ACT1-CUP1 reporters harboring substitutions at the SS. The most significant decreases in Cu^2+^ tolerance were observed using reporters containing the A3C substitution at the 5′ SS, substitutions of the branch point adenosine (A259C or A259G), or substitutions flanking the branch point (U257C and C260G) (**Fig. 4B**). The large impact of *ecm2Δ* on A3C reporter splicing was intriguing since this substitution is only limiting for the 2^nd^ catalytic step (Liu et al. 2007; Eysmont et al. 2019). Primer extension analysis of 1^st^- and 2^nd^-step splicing products confirmed a strong defect in exon ligation for the A3C reporter in the absence of Ecm2 (**Supplemental Fig. S1**). Ecm2 can therefore have opposing effects on the 2^nd^-step active site: it can inhibit its formation but also promote 2^nd^-step catalysis on a substrate containing the A3C 5′ SS substitution. It is possible that Ecm2 has distinct functions in both spliceosome structural transitions and in each catalytic reaction.

Deletion of ECM2 improved Cu^2+^ tolerance of yeast containing the A3U or G5A 5′ SS reporters (**Fig. 4B**). However, analysis of 1^st^- and 2^nd^-step splicing products showed similar splicing efficiencies in the presence or absence of Ecm2 (**Supplemental Fig. S1**). We did not study how decay of unspliced RNAs or splicing intermediates influenced these primer extension results (Liu et al. 2007). Therefore, it is difficult to conclude from the primer extension assay if *ecm2Δ* truly changed the splicing efficiencies for the A3U and G5A substrates.

Nonetheless, the increase in Cu^2+^ tolerance observed with the G5A mutant in *ecm2Δ* yeast was noteworthy since this substitution can result in use of a cryptic, upstream 5′ SS (Parker and Guthrie 1985; Lesser and Guthrie 1993a; Kandels-Lewis and Séraphin 1993). We used a modified ACT1-CUP1 reporter with competing 5′ SS to test whether or not Ecm2 changes cryptic SS usage (**Fig. 4C**, **Supplemental Fig. S2**). When Ecm2 was present in the yeast, we detected usage of both the cryptic (21±1% of spliced products) and normal 5′ SS. However, in the absence of Ecm2 use of the cryptic 5′ SS was greatly reduced (7±4% of spliced products, **Fig. 4D**). This represents at least a 3-fold decrease based on our lower limit of detection. Combined, our results demonstrate that Ecm2 impacts the spliceosome active site to alter splicing of pre-mRNAs with non-consensus SS and permit the usage of an alternate, cryptic 5′ SS.

### Genetic Interactions Between Ecm2 and the U6 snRNA 1^st^- and 2^nd^-Step Alleles

Like *ecm2Δ*, a number of alleles of the U6 snRNA and Prp8 suppress Prp16 ATPase mutations, have limited impact on splicing of ACT1-CUP1 reporters harboring consensus SS, and can promote or block splicing of reporters with particular SS substitutions (Eysmont et al. 2019; Liu et al. 2007; Query and Konarska 2004; Mayerle et al. 2017; McPheeters 1996). Many of these mutants have been categorized as 1^st^-or 2^nd^-step alleles since, in addition to causing *ts* or *cs* phenotypes, they promote one of the catalytic steps of splicing over the other (**Fig. 5A**). Since *ecm2Δ* and 1^st^- and 2^nd^-step alleles share common features, we tested for genetic interactions between these alleles and *ecm2Δ*. We first examined interactions with the U6 snRNA U57C and U57A mutations which promote the 1^st^ and 2^nd^ steps, respectively (McPheeters 1996; Liu et al. 2007).

**Figure 5.**
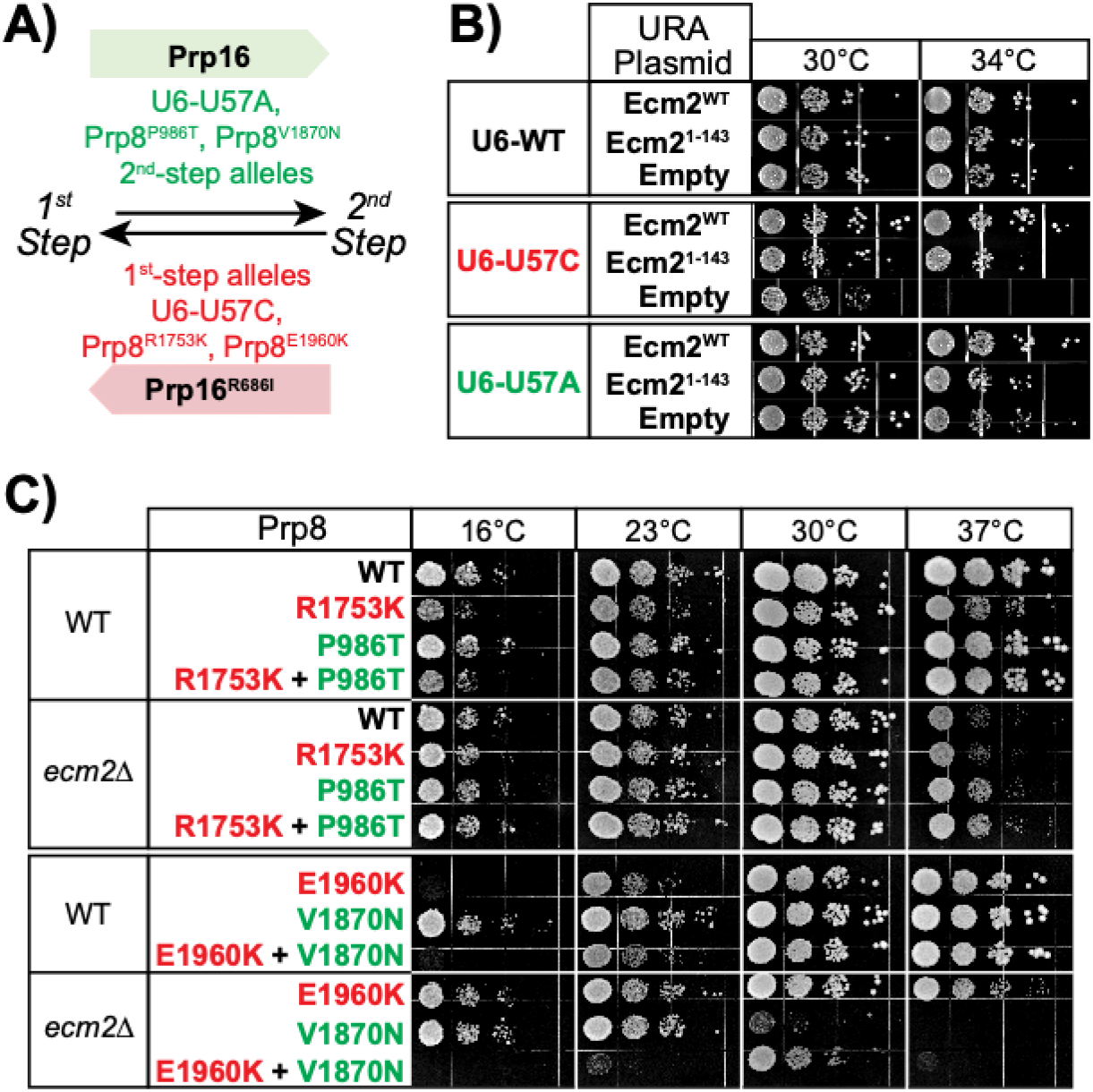
Genetic Interactions between Ecm2 and U6 or Prp8 1^st^- and 2^nd^-Step Alleles. (**A**) Illustration of how alleles of Prp8, Prp16, and U6 function to promote the 1^st^ or 2^nd^ step of splicing. (**B**) A 1^st^-or 2^nd^-step allele of U6 (red and green, respectively) was combined with URA3 plasmids either lacking or coding for Ecm2 variants in *ecm2Δ* yeast. The strains were then tested for suppression or exacerbation of temperature-dependent growth phenotypes. Yeast were plated on-URA dropout media and imaged after 2 days of growth. (**C**) 1^st^- and 2^nd^-step alleles of Prp8 (red and green, respectively) were combined with *ecm2Δ* and tested for suppression or exacerbation of temperature-dependent growth phenotypes. Yeast were plated on YPD media and imaged after 3 (23, 30, or 37°C) or 10 (16°C) days of growth.

The U6-U57A mutation had no effect on yeast growth at 16, 23, or 30°C in the absence of Ecm2 or in the presence of Ecm2^WT^ or Ecm2^1-143^ (which contains only the U6-binding domain, **Fig. 5B**and data not shown). The U6-U57C mutation also had no impact on growth at 16 or 23°C but was slower growing at 30°C. U6-U57C yeast containing only the empty URA3 plasmid displayed a slower-growing phenotype compared to those containing plasmids for Ecm2^WT^ or Ecm2^1-143^.

Strains deleted of both Ecm2 and U6 (*ecm2Δ snr6Δ*) failed to grow at 37°C even when they contained plasmids encoding for WT U6 and Ecm2 (data not shown). However, we were able to assay growth at 34°C. When yeast contained the 1^st^-step, U6-U57C allele, we observed a strong synthetic sick interaction with *ecm2*Δ that was partially rescued by expression of Ecm2^WT^ or Ecm2^1-143^, with the former producing stronger rescue than the latter. In contrast, we observed only a weak synthetic genetic interaction between *ecm2Δ* and the 2^nd^-step allele, U6-U57A (**Fig. 5B**).

The interactions of *ecm2Δ* with these U6 mutants are most similar to those of 1^st^-step alleles in other splicing factors like Prp8 (Liu et al. 2007). When U6 mutations are present, loss of Ecm2 promotes the 1^st^ step of splicing and presence of Ecm2 promotes the 2^nd^ step. These results differ from those obtained upon deletion of the non-essential factor Isy1 (**Fig. 1C**): *isy1Δ* is synthetic lethal with U57A and suppresses the *ts* phenotype of U57C (Villa and Guthrie 2005). Thus, Isy1 appears to act as a 1^st^-step splicing factor when U6 is mutated, consistent with Isy1 release prior to the 2^nd^ step (Plaschka et al. 2019), while Ecm2 can act as a 2^nd^-step factor and is consistent with its presence throughout both catalytic stages of splicing (**Fig. 1A**).

### Genetic Interactions Between Ecm2 and Prp8 1^st^- and 2^nd^-Step Alleles

Genetic interactions between *ecm2Δ* and Prp2, Prp16, and U2 stem II and Ecm2-control of 5′ SS usage support a role for Ecm2 in the 1^st^ step of splicing. However, genetic interactions with U6-U57 mutants and results using the A3C splicing reporter support an additional role for Ecm2 in the 2^nd^ step. We next tested if *ecm2Δ* would show genetic interactions with 1^st^-or 2^nd^-step alleles of Prp8 or both. We individually combined *ecm2Δ* with two 1^st^-step alleles of Prp8 (Prp8^R1753K^ or Prp8^E1960K^) or two 2^nd^-step alleles (Prp8^P986T^ or Prp8^V1870N^). For the 1^st^-step alleles, deletion of Ecm2 weakly suppressed the *cs* phenotype of Prp8^R1753K^ and strongly suppressed the *cs* phenotype of Prp8^E1960K^ (**Fig. 5C**). Neither Prp8 1^st^-step allele was able to completely correct the *ts* phenotype of *ecm2Δ*; although, slightly improved growth was observed at 37°C for yeast containing Prp8^E1960K^ (**Fig. 5C**).

When *ecm2Δ* was combined with the 2^nd^-step alleles, we observed slightly improved growth at 37°C for yeast containing Prp8^P986T^. A stronger genetic interaction was observed with the Prp8^V1870N^. This 2^nd^-step allele exacerbated the *ts* phenotype of *ecm2Δ*, causing a strong growth defect at 30°C and no growth at 37°C (**Fig. 5C**). The growth defect of Prp8^V1870N^ was partially corrected at 30°C by combining 1^st^ and 2^nd^ Prp8 alleles (Prp8^V1870N,E1960K^). However, this also resulted in stronger growth defects at other temperatures.

The *cs* suppression we observe of the Prp8^E1960K^ 1^st^-step allele and *ts* exacerbation of the Prp8^V1870N^ 2^nd^-step allele are consistent with *ecm2Δ* acting as a 2^nd^-step allele and facilitating exit of the spliceosome from the 1^st^-step catalytic conformation. Both the Prp8^E1960K^ and Prp8^V1870N^ substitutions are located within Prp8’s RNaseH domain. Like U2 stem II, the RNaseH domain is both highly dynamic and toggles between alternate structures (Mayerle et al. 2017; Schellenberg et al. 2013). Ecm2 may impact Prp8-RNaseH dynamics or vice versa to support the 1^st^-step reaction. This is in juxtaposition to the results obtained with the U6 mutants, which were consistent with Ecm2 having a role in the 2^nd^ step.

## DISCUSSION

Using genetic and biochemical assays of splicing in cells, we have shown that yeast Ecm2 impacts multiple steps during the catalytic phase of splicing and that loss of Ecm2 perturbs how the spliceosome processes pre-mRNAs containing non-consensus SS. Deletion of *ECM2* results in genetic interactions with several structural switches in the spliceosome including U2 stem II, the RNaseH domain of Prp8, and the ATPases that control entry to and exit from the 1^st^ step (Prp2 and Prp16, respectively). In sum, our data show that Ecm2 plays significant roles in spliceosome catalysis in addition to a function during activation (Xu and Friesen 2001).

### Ecm2 Modulates the Catalytic Steps of Splicing

Our results support functions for Ecm2 during both catalytic steps in splicing. The differing genetic interactions we observe between *ecm2Δ* and U6 or Prp8 mutants suggest a more complicated mechanism from that of other alleles that exhibit more consistent genetic interactions (for example, a 2^nd^-step allele of *cef1* suppresses phenotypes of both 1^st^-step *prp8* and U6 alleles) (Query and Konarska 2012). Our results could be explained by distinct and genetically separable functions for Ecm2 during the 1^st^ and 2^nd^ catalytic steps with Prp8 mutations revealing a role in the former and U6 mutations revealing a role in the latter. Since Ecm2 has only been observed to make contact with U2 stem IIb in 1^st^-step cryo-EM structures, it is possible that this interaction contributes to Ecm2’s distinct functions during each catalytic step.

Several of our observations with Ecm2 are similar to those previously reported for Cwc2 and Isy1 (Hogg et al. 2014; Villa and Guthrie 2005; Rasche et al. 2012). Cwc2, Ecm2, and Isy1 form a highly interconnected network of interactions with one another, the U6 snRNA, the intron, and a number of other splicing factors (**Figs. 1C**, **6A**) (Galej et al. 2016; Wan et al. 2016). All three proteins can suppress Prp16 mutations and have synthetic lethal interactions with active site U2/U6 helix Ia or Ib mutations (Hogg et al. 2014; Villa and Guthrie 2005; Xu et al. 1998). Neither Cwc2, Isy1, nor Ecm2 is essential for yeast growth, and cells remain viable even when Cwc2 and Ecm2 are both absent albeit with a significant *ts* growth defect (Hogg et al. 2014; Villa and Guthrie 2005; Xu and Friesen 2001). In addition, loss of Ecm2, loss of Isy1, or mutation of Cwc2 results in specific splicing defects in reporter pre-mRNAs with non-consensus SS (**Fig. 4B**) (Hogg et al. 2014; Villa and Guthrie 2005). This implies that spliceosomes missing one of these factors possess different substrate preferences and fidelity phenotypes. This is intriguing since *ecm2Δ* only results in a growth defect at high temperatures and splicing of pre-mRNAs containing consensus SS remains efficient (**Fig. 4B**). Thus, it is possible that yeast could bias the splicing of particular pre-mRNAs with non-consensus SS by regulating the Cwc2, Ecm2, and/or Isy1 content of spliceosomes without significantly compromising cellular splicing efficiency. This possibility has previously been proposed by Villa and Guthrie, who noted that deletion of Isy1 results in reduced fidelity of 3′ SS selection (Villa and Guthrie 2005).

**Figure 6.**
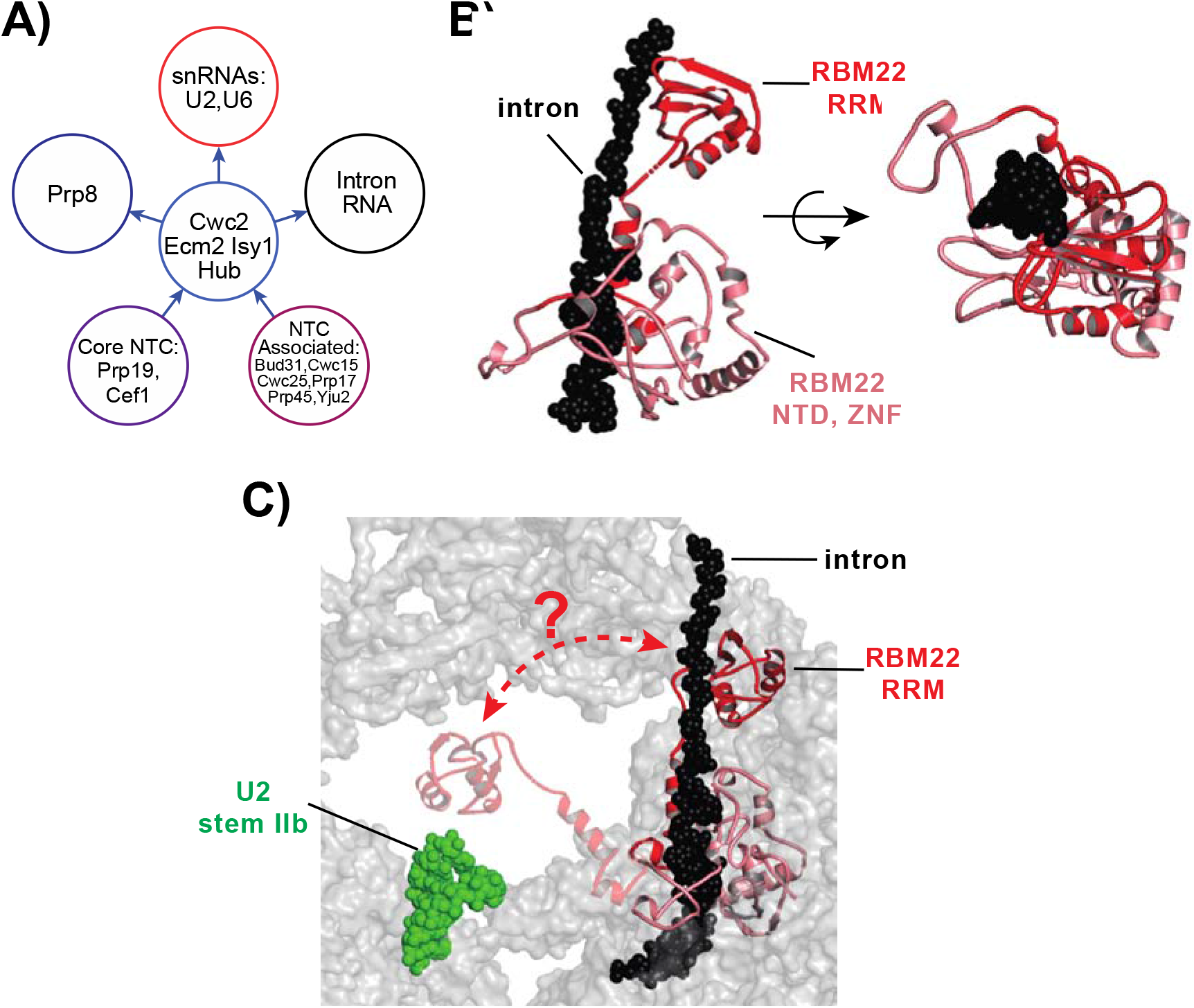
The Cwc2/Ecm2/Isy1 Interaction Network and Structure of Human RBM22. (**A**) A large number of splicing factors interact with Cwc2, Ecm2, and/or Isy1 suggesting that these proteins form a network hub for modulating spliceosome activity. In this model, regulatory signals could flow into the hub from the NTC and NTC-related proteins and outwards to the spliceosome active site consisting of the intron, Prp8, and U2/U6 snRNAs. (**B**) Two views of the cryo-EM structure of RBM22 from a human C complex spliceosome. Domains of RBM22 are noted and intronic RNA downstream of the 5′ SS is shown in black spacefill. Note that RBM22 completely encircles the RNA. Structure from 6EXN.pdb. (**C**) Hypothetical model for formation of the structure shown in panel (B). The RRM domain of RBM22 could make transient contact with human U2 stem IIb to allow for docking of the intron and subsequent wrapping. Structures in panels (B) and (C) are from 5YZG.pdb. The hypothetical model in panel (C) was created using PyMol.

While there is some overlap in how Isy1 or Ecm2 loss or Cwc2 mutation impact splicing of non-consensus SS, the proteins also exert unique influences of the spliceosome. For example, *isy1Δ* and the Cwc2^F183D^ mutant improve splicing of reporter pre-mRNAs containing a A302U 3′ SS, but *ecm2Δ* only minimally changes A302U splicing (**Fig. 4B**). In addition, *ecm2Δ* changes Cu^2+^ tolerance with the G5A reporter but this is unaffected by *isy1Δ* or Cwc2^F183D^ (Villa and Guthrie 2005; Hogg et al. 2014). Cellular splicing could thus be optimized for specific SS by independently controlling Cwc2, Ecm2, and Isy1 stoichiometry with spliceosomes.

These factors might also impact mRNA isoform production since Ecm2 additionally permits usage of an alternative 5′ SS (**Fig. 4D**). Interestingly, the non-essential yeast splicing factor Bud31 is required for use of an alternative 5′ SS in the SRC1 mRNA (Saha et al. 2012), and Bud31 directly contacts the U6 snRNA, Ecm2, and Cwc2 in the yeast spliceosome (Plaschka et al. 2019). Bud31 and Ecm2 could permit promiscuous 5′ SS use by similar mechanisms, although this has not yet been studied. In summary, spliceosomes may be fine-tuned by the presence or absence of non-essential splicing factors like Ecm2, and currently little is known about how the compositions of spliceosomes vary inside cells.

### Consequences of a Dynamic Ecm2/U2 Stem II Interaction During Splicing

The spliceosome contains a number of proposed switches in which components toggle between one conformation or another at different stages of the reaction (Abelson 2017). The U2 snRNA contains several of these switches including a U2 stem IIa-to-IIc conformational change during activation (van der Feltz and Hoskins 2019). In addition, it has also been proposed that stem IIc switches transiently back to stem IIa between the catalytic steps of splicing before re-forming IIc during the 2^nd^ step (Perriman and Ares 2007; Hilliker et al. 2007). This mechanism was based in part on the observation that stem II substitutions that destabilize stem IIc (or stabilize stem IIa) can suppress *cs* alleles of Prp16. Our observations that *ecm2Δ* also suppresses Prp16 *cs* alleles (**Fig. 2**) and Ecm2 contacts U2 stem IIb in C complex may provide an alternate explanation.

We propose that Prp16 indirectly disrupts the Ecm2/stem II interaction during remodeling of the spliceosome between the 1^st^ and 2^nd^ steps. Eliminating or weakening this interaction by stem II mutation can suppress Prp16 *cs* alleles by destabilizing the 1^st^-step conformation. This explanation is supported by cryo-EM structural data in which a transient contact between U2 stem IIb/c and the C-terminal RRM of Ecm2 is observed in 1^st^-step complexes (B* and C complexes) but not those preceding or following (B^act^ and C* complexes) (Rauhut et al. 2016; Galej et al. 2016; Fica et al. 2017; Yan et al. 2016a; Wan et al. 2016, 2019). While additional structural information is needed for the on-pathway intermediates during 2^nd^-step active site assembly, stem IIa has not yet been observed in C* spliceosomes and accommodation of stem IIa in these complexes may be incompatible with binding of splicing factors (Prp17) and U2 snRNP interactions with Syf1 (Fica et al. 2019; Wan et al. 2018; Liu et al. 2017; Fica et al. 2017; Yan et al. 2016b). In light of these observations, stem IIc could remain intact throughout catalysis, and IIc-to-IIa toggling occurs later during spliceosome disassembly or U2 snRNP reassembly (Rodgers et al. 2015; Yan et al. 1998). Regardless, further work is needed to characterize the short-lived intermediates that form during the 1^st^- to 2^nd^-step transition.

The viability of *ecm2Δ* and stem IIbΔ strains (**Fig. 3B**) (Xu and Friesen 2001; Ares and Igel 1990) show that the Ecm2/stem II interaction is not essential for yeast splicing. It is notable, however, that *ecm2Δ* exhibits synthetic lethal interactions with multiple stem II mutations. This includes, to our knowledge, the first genetic data showing synthetic lethality with the U2-2,4 mutant, which stabilizes stem IIa. This supports the notion that stem IIa must be disrupted during splicing and complements ample evidence for the importance of stem IIc formation (Perriman and Ares 2007; Hilliker et al. 2007). It is possible that the Ecm2/stem II interaction only becomes limiting for splicing when stem IIa/c toggling is disturbed or when the active site is destabilized by SS mutations.

### Implications for Human RBM22 and Wrapped Intron Formation

The evolutionary histories of Cwc2, Ecm2, and RBM22 have not been studied, and it is uncertain how RBM22 may have evolved to functionally replace both proteins. Based on sequence alignments and crosslinking studies, it has been proposed that Cwc2 and RBM22 share a common function in binding U6 and interacting with the spliceosome’s catalytic elements (Rasche et al. 2012). However, when fragments of the yeast and human C complex spliceosomes are aligned, RBM22 most closely mimics the interactions of Ecm2 with the U6 snRNA (**Supplemental Movie S1)**. In terms of U6 interaction, we believe that RBM22 and Ecm2, not Cwc2, are closer structural homologs.

Both RBM22 and Cwc2 bind the intronic RNA just downstream of the 5′ SS. The RBM22/intron interaction contains an unusual and distinctive structure not observed with Cwc2. In human C and P complex spliceosomes, RBM22 completely encircles the intron (**Figure 6B)**(Fica et al. 2019; Zhan et al. 2018). It is unlikely that this wrapped intron structure would form by threading of the intron through RBM22. Insights from Ecm2 provide a plausible mechanism for its formation. The C-terminal RRM of RBM22 could transiently interact with U2 stem IIb/c—analogous to the interaction between Ecm2 and stem II in yeast (**Figure 6C**). This could open RBM22 for intron binding and enable subsequent wrapping of the intron after disruption of the RRM/stem II interaction.

Analyses of cryo-EM structures reveal that movement of RBM22 towards U2 stem II is not occluded by presence of other splicing factors and stem IIb is within an accessible distance for the RRM, assuming structural flexibility of the linker between the ZNF and RRM domains. There is some biochemical evidence for a RBM22/U2 snRNA interaction: anti-RBM22 antibodies can immunoprecipitate (IP) small amounts of the U2 snRNA from C complex spliceosomes after proteinase K treatment and without co-IP of the U5 or U6 snRNAs—consistent with a direct interaction (Rasche et al. 2012). If a transient RBM22/U2 interaction is necessary for intron wrapping, U2 snRNA stem II may thus act as a chaperone for formation of this protein/RNA complex.

## MATERIALS AND METHODS

Yeast strains and plasmids used in these studies are described in **Supplemental Tables S1** and **S2**. Yeast transformation and growth were carried out using standard techniques and media.

### Genetic Deletions of ECM2

Deletion of the ECM2 gene was carried out by replacement of the gene with an antibiotic resistance cassette (hygromycin or nourseothricin) by homologous recombination in the appropriate parental strain (**Supplemental Table S1**, (Goldstein and McCusker 1999)).

### Cloning of ECM2 and Site-Directed Mutagenesis

ECM2 along with 300 base pairs of up- and downstream DNA was amplified from yeast genomic DNA by PCR. The resulting product was digested with NotI and SalI restriction enzymes and then ligated into pRS416 (URA3 CEN6) at those same restriction sites to create plasmid pAAH1056 containing the WT ECM2 gene. Novel mutants of Ecm2 were generated using inverse polymerase chain reaction (PCR) with Phusion DNA polymerase (New England Biolabs; Ipswich, MA). All plasmids were confirmed by sequencing.

### *ACT1-CUP1* Copper Tolerance Assays

Yeast strains expressing *ACT1-CUP1* reporters were grown to mid-log phase in-leu-trp dropout media to maintain selection for the plasmids, adjusted to OD_600_ = 0.5 and equal volumes were spotted onto plates containing 0-2.5 mM CuSO_4_ (Lesser and Guthrie 1993b; Carrocci et al. 2018). Plates were scored and imaged after 3 days growth at 30°C.

### Temperature Growth Assays

Yeast strains were grown to mid-log phase at permissive temperatures in YPD or-ura dropout liquid media. Cell growth was then quantified by measuring OD_600_. Equal volumes of cells were diluted to an OD_600_ = 0.5 were stamped onto YPD-agar plates and incubated at 23°C, 30°C or 37°C for three days or at 16°C for ten days before imaging.

### Primer Extension

Cells were grown at 30°C in 25 mL yeast-leu-trp dropout liquid media until OD_600_ reached 0.5–0.8, and 10 OD_600_ units were collected by centrifugation. Total yeast RNA was isolated following the MasterPure™ Yeast RNA Purification Kit (Epicenter, Madison, WI) protocol with minor changes as previously described (Carrocci et al. 2017). Primer extension was performed with IR dye conjugated probes yAC6: /5IRD700/GGCACTCATGACCTTC and yU6: /5IRD700/GAACTGCTGATCATCTCTG. purchased from Integrated DNA Technologies (Skokie, IL USA) (Carrocci et al. 2017; Kaur et al. 2020). Gels were imaged with the Amersham IR Typhoon 5 (GE Healthcare) excitation at 685 nm, emission filter 720BP20, PMT voltage of 700V, and 100 μm pixel size. Band intensities were quantified with ImageQuant TL v8.1 (GE Healthcare).

### Structural Alignments and Figure Creation

Structural alignments of portions of human and yeast spliceosome complexes were carried out using PyMol by aligning to the U6 snRNA. Aligned structures of yeast spliceosomes were obtained from PyMOL4Spliceosome (https://github.com/mmagnus/PyMOL4Spliceosome) (Magnus et al. 2019). Figures and movies containing molecular structures were generated using Pymol (Schrödinger).

## SUPPLEMENTAL MATERIAL

Supplemental material is available for this article.

## ACKNOWLEDGEMENTS

We thank Maggie Rodgers and Brexton Turner for carrying out initial experiments on Ecm2. We thank Manny Ares (UC Santa Cruz), Charles Query (Albert Einstein College of Medicine), David Brow (U. Wisconsin-Madison), Christine Guthrie (UCSF), and Magda Konarska (U. Warsaw) for providing strains and plasmids used in these studies. We thank Eric Montemayor (UW-Madison) for help in creating Figure 6. We thank Tucker Carrocci (Yale U.), Sarah Hansen (U. Utah), Karli Lipinski (UW-Madison), and Maggie Rodgers (Johns Hopkins U.) for critical reading of the manuscript.

## FUNDING

This work was supported by the National Institutes of Health (R01 GM112735 and R35 GM136261 to AAH); a Shaw Scientist Award (AAH); and Hilldale Undergraduate Research Scholarships (BN and CS).

